# Detection and characterization of low and high genome coverage regions using an efficient running median and a double threshold approach

**DOI:** 10.1101/092478

**Authors:** Dimitri Desvillechabrol, Christiane Bouchier, Sean Kennedy, Thomas Cokelaer

## Abstract

**Motivation:** Next Generation Sequencing (NGS) provides researchers with powerful tools to investigate both prokaryotic and eukaryotic genetics. An accurate assessment of reads mapped to a specific genome consists of inspecting the *genome coverage* as number of reads mapped to a specific genome location. Most current methods use the average of the genome coverage (*sequencing depth*) to summarize the overall coverage. This metric quickly assess the sequencing quality but ignores valuable biological information like the presence of repetitive regions or deleted genes. The detection of such information may be challenging due to a wide spectrum of heterogeneous coverage regions, a mixture of underlying models or the presence of a non-constant trend along the genome. Using robust statistics to systematically identify genomic regions with unusual coverage is needed to characterize these regions more precisely.

**Results:** We implemented an efficient running median algorithm to estimate the genome coverage trend. The distribution of the normalized genome coverage is then estimated using a Gaussian mixture model. A `z`-score statistics is then assigned to each base position and used to separate the central distribution from the regions of interest (ROI) (i.e., under and over-covered regions). Finally, a double threshold mechanism is used to cluster the genomic ROIs. HTML reports provide a summary with interactive visual representations of the genomic ROIs.

**Availability:** An implementation of the genome coverage characterization is available within the `Sequana` project. The standalone application is called `sequana_coverage`. The source code is available on `GitHub` (http://github.com/sequana/sequana), and documentation on `ReadTheDocs` (http://sequana.readtheodcs.org). An example of HTML report is provided on http://sequana.github.io.

## 1 Introduction

Sequencing technologies allow researchers to investigate a wide range of genomic questions [Goodwin et al., 2016], covering research fields such as the expression of genes (transcriptomics) [Wang et al., 2009], the discovery of somatic mutations, or the sequencing of complete genomes of cancer samples to name a few examples [Meyerson et al., 2010, Iorio et al., 2016]. The emergence of the second generation sequencing, which is also known as Next-Generation Sequencing or NGS hereafter, has dramatically reduced the sequencing cost. This breakthrough multiplied the number of genomic analyses undertaken by research laboratories but also yielded vast amount of data. Consequently, NGS analysis pipelines require efficient algorithms and scalable visualization tools to process this data and to interpret the results.

Raw data generated by NGS experiments are usually stored in the form of *sequencing reads* (hereafter simply called reads). A read stores the information about a DNA fragment and also an error probability vector for each base. Read lengths vary from 35-300 bases for current short-read approaches [Goodwin et al., 2016] to several tens of thousands of bases possible with long-read technologies such as Pacific Biosciences [Eid et al., 2009, Lee et al., 2014] or Oxford Nanopore [Eisenstein, 2012].

After trimming steps (quality, adapter removal), most NGS experiments will require mapping the reads onto a genome of reference [Li, 2013]. If no reference is available, a *de-novo* genome assembly can be performed [Bankevich et al., 2012]. In both cases, reads can be mapped back on the reference taking into account their quality. We define the *genome coverage* as the number of reads mapped to a specific position within the reference genome. The theoretical distribution of the genome coverage has been thoroughly studied following the seminal work of Lander-Waterman model [Lander and Waterman, 1988, Wendl et al., 2005]. A common metric used to characterize the genome coverage is the *sequencing depth:* the empirical average of the genome coverage. Its unit is denoted X. An example of a genome coverage with a sequencing depth of about 450 X is shown in Fig.1.

**Fig. 1.**
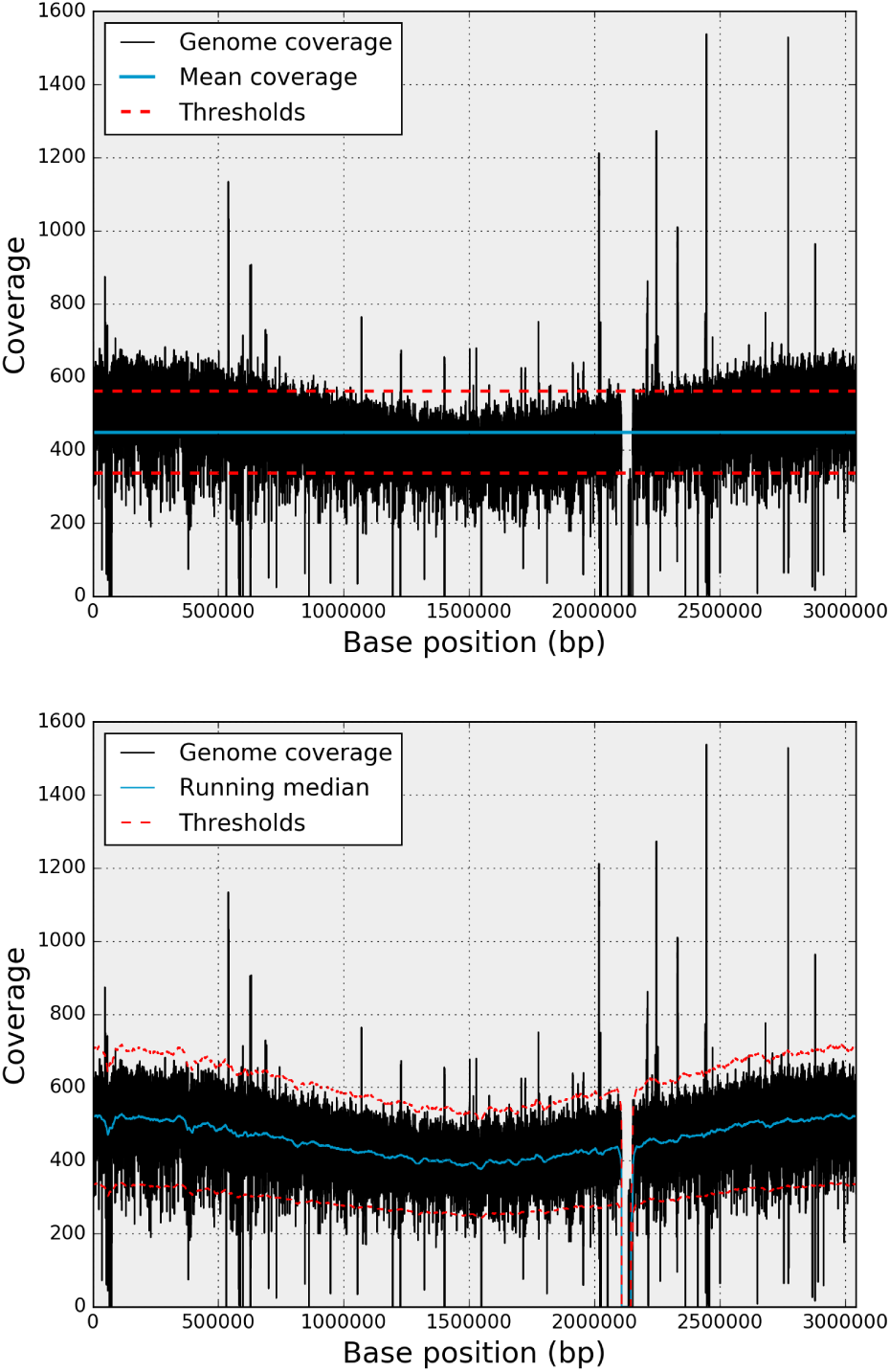
Example of a genome coverage series (in black in both panels). The genome coverage corresponds to the bacteria test case (see text). It contains a deleted region (around 2.2 Mbp) and various under and over-covered regions (from 100 bp to several Kbp). Although the sequencing depth is about 500 X, there is non-linear trend from 500 X on both ends to 400 X in the middle of the genome. The top panel shows the sequencing depth (blue horizontal line) and two arbitrary fixed thresholds (dashed red lines) at 400 X and 500 X. Due to the non-linear trend, the fixed thresholds lead to an increase of Type I and Type II errors. On the contrary, in the bottom figure, the trend is estimated using a running median (red line) and adaptive lower and upper thresholds (dashed red lines) can be derived.

The required sequencing depth depends on the experimental application. For instance, to detect human genome mutations, singlenucleotide polymorphisms (SNPs), and rearrangements, a 50 X depth is recommended [Goodwin et al., 2016] to be able to distinguish between sequencing errors and true SNPs. In contrast, the detection of rarely expressed genes in transcriptomics experiments often requires greater sequencing depth. However, greater sequencing depth is not always desirable. Indeed, in addition to a higher cost, ultra-deep sequencing (large sequencing depth in excess of 1000 X) may be an issue for *de-novo* genome assembly [Mirebrahim et al., 2015]

The Lander-Waterman model also provides a good theoretical estimate of the required redundancy to guarantee that for instance all nucleotides are covered at least *N* times. This is, however, a theoretical estimate that does not take into account technological and biological limitations: some regions are indeed difficult to efficiently map (e.g., repetitive DNA). Furthermore, the genome coverage may also contain a non-constant trend or additional sequence not present in the reference. The genome coverage example in Fig. 1 demonstrates these different features.

While the sequencing depth metric provides a quick understanding about the quality of the mapping, the genome coverage can be further used to identify regions that are significantly under- or over-covered. Hereafter, these regions of interest (ROI) are denoted low-ROIs and high-ROIs, respectively.

In order to detect low and high-ROIs, a simple and fast approach consists in setting two arbitrary thresholds bounding the sequencing depth. There are two major drawbacks with this approach. First, as shown in Fig. 1A, with a fixed threshold, one may detect numerous false signals (type I errors) or fail to detect real events (type II errors). Secondly, the threshold is fixed manually and lacks a robust statistics. An alternative is to estimate the genome coverage profile histogram [Linder et al., 2013] from which a `z`-score statistics can be used to identify outliers more precisely. Yet, since the genome coverage may contain low and high frequency fluctuations, the statistics will also suffer from Type I and II errors.

In this paper, we describe an approach that first estimates the genome coverage trend using a running median in place of a running mean. It can be employed to normalize the genome coverage vector and calculate a robust statistic (`z`-score) for each base position. This allows us to obtain robust low and high thresholds at each base position.

In section 2, we describe the data sets used throughout the paper as test-case examples. In section 3, we describe (i) the running median used to detrend the genome coverage and (ii) the statistical method used to characterize the central distribution from which outliers can be identified and (iii) we propose a double threshold method to cluster ROIs. Finally, in section4, we describe the standalone application, *sequana_coverage,* and its potential applications for NGS research projects.

## 2 Material

Three test-cases of genome coverage are presented here, covering representative organisms and sequencing depths. The genome coverage data sets are in BED (Browser Extensible Data) format, a tabulated file containing the coverage, reference (e.g. chromosome number, contig) and position on the reference. BED files can be created using `bedtools` (http://bedtools.readthedocs.io), in particular the `genomecov tool`.

We first considered a bacteria from a study about methicillin resistant *Staphilococcus aureus* [Tong et al., 2015]. One circular chromosome of 3 Mbp is present. The sequencing depth is 450 X and the genome coverage exhibits a non-constant trend along the genome (see Fig.1). This pattern, often observed in rapidly growing bacteria, is the result of an unsynchronized population where genome replication occurs bidirectionally from a single origin of replication [Bremer et al., 1977, Prescott et al., 1972]. The proportion of outliers (see Sec 3.2 for a formal definition) is about 2.5 % of the total bases. The original data sets (Illumina sequencing reads, paired-end, 100bp) are available at the European Nucleotide Archive (ENA; http://www.ebi.ac.uk/ena/) under study accession number ERP000130 (ERR036019). The reference is FN433596. A genome coverage in BED format is available on Synapse (https://www.synapse.org) under number syn7211116.

The second organism is a virus with a sequencing depth of 1000 X [Combredet et al., 2003]. The circular plasmid sequencing, which contains the virus chromosome, is 19 795 bp-long. About 14 % of the genome coverage contains large or low coverage regions (outliers). It also contains two large under-covered regions (one partially under-covered and one not covered at all) as shown in Fig. 2. A genome coverage BED file is available on Synapse under number syn7211115. The reference can be found on ENA website (http://www.ebi.ac.uk/ena/data/view/JB409847).

The third test case is the fungus *(Schizosaccharomyces pombe)* [Wood et al., 2002]. The genome coverage has a sequencing depth of 105 X. It has three non-circular chromosomes of 5.5 Mbp, 4.5 Mbp and 2.5 Mbp. The references from ENA are CU329670.1, CU329671.1 and CU329672.1 (X54421.1). Although we will look at the first chromosome only (1% of outliers), the tools presented hereafter handles circular chromosomes and multiple chromosomes. A genome coverage file in form of a BED file is available on Synapse (syn7561732).

**Fig. 2.**
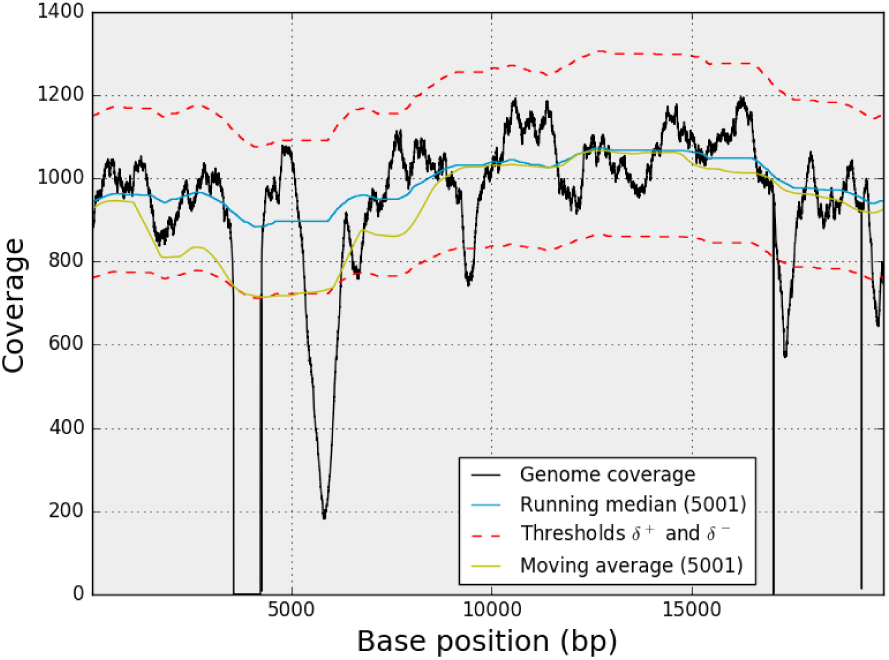
Comparison of the running median and moving average estimators (virus case). The sequencing depth is 930 X and the genome coverage has a deleted region situated around b = 4 000 as well as an under-covered region at b =6 000. The moving average is less robust to outliers or deleted regions. For instance, the region around b = 5000 is biased due to the presence of a deleted region, which increases the rate of false alarms.

## 3 Methods

### 3.1 Detrending the genome coverage

The genome coverage function is denoted *C(b)* where *b* is the base (nucleotide) position on the genome of reference. The genome coverage and reference lengths are denoted *N*. For simplicity, we drop the parentheses and refer to the genome coverage as *C_b_*. The empirical sequencing depth (average of genome coverage) is denoted δ =*C̅_b_*. Ideally, *C_b_* is made of a continuous homogeneous central region. In practice, however, this may be interrupted by a succession of under- and over-covered regions: the genomic ROIs that we want to detect.

A naive classifier consists in setting two fixed thresholds *δ^−^* and *δ^+^* whereby low and high ROIs are defined as 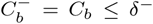 and 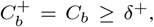 respectively. If 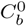 denotes the remaining data such that 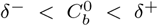, then the genome coverage can be written as 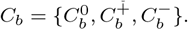

The advantage of the fixed-thresholds method is that it is conceptually simple and computationally inexpensive. However, there are two major drawbacks manifest. First, as shown in Fig.1-A, false negatives and false positives will increase as soon as there is a non-constant trend present in the data. It may be a low frequency trend as shown here but high frequency trend are also present (see e.g., Fig 2). Also of importance is that an arbitrary choice of threshold(s) is unsatisfactory from a statistical point of view since we cannot associate any level of significance to a genomic region.

In order to account for a possible trend in the genome coverage series (and remove it), a standard method consists in dividing the series by a representative alternative such as its moving average or running median.

The moving average (MA) is computed at each position, *b*, as the average of W data points around that position and defined as follows:

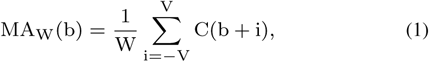

where *W* is the length of the moving window (odd number) and *V* = (*W* – 1)/2. Note that the first and last *V* values are undefined. However, in the case of circular DNA (e.g., virus case), then the first and last V points are defined since *C_b_* is now a circular series.

Similarly, the running median (RM) is computed at each position, *b*, as the median of W data points around that position:

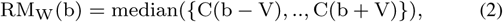

where *W* and *V* are defined as before and the median function is defined as the middle point of the sample set (half of the data is below the median and half is above). A mathematical expression of the median and running median are given in the Appendix section (Eq. 8).

The mean estimator is commonly used to estimate the central tendency of a sample, nevertheless it should be avoided in the presence of extraneous outliers, which are common in NGS genome coverage series (see e.g., Fig. 1). Fig. 2 shows the impact of outliers when using a moving average or a running mean. We will use the running median only and define the normalized genome coverage as follows:

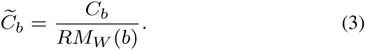

We will use the tilde symbol for all metrics associated with the normalized genome coverage,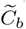. For instance, 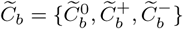.

The running median is used in various research fields, in particular in spectral analysis [Percival, 1993] to estimate the noise floor while ignoring biases due to narrow frequency bands (e.g., [Balasubramanian et al., 2005]). Here, the goal is to avoid narrow peaks but also to be insensitive to long deleted regions. This can be a major issue in NGS as the running median estimator complexity is a function of the window length. Indeed the running median algorithm involves the sorting of a sample of length W at each position of the genome. So, the running median estimator must be efficient and scalable. This is not an issue in spectral analysis and most fields where running median are used but is a bottleneck for NGS analysis. As explained in the Appendix section, the complexity of the sorting part is in *Ο*(*n*^2^) in the worst case but similarly to the moving average, one can take advantage of the rolling window and the fact that the previous block is already sorted. We report a Python implementation with a complexity as low as *Ο*(log *n*) (see Appendix). Briefly, it takes advantage of (i) a bisection method to insert new elements inside an already sorted block of data and (ii) an efficient Python data structure optimised to insert and delete elements in a list. In our implementation, both the moving average and running median have the ability to account for circular DNA data, which is essential to handle circular series.

If we normalize the genome coverage from the bacteria example (Fig.1), we obtain the results shown in Fig.3. Finally, note that the genome coverage being discrete, the running median is also discrete as well as the normalized genome coverage. The discreteness will become more pronounced as sequencing depth decreases.

**Fig. 3.**
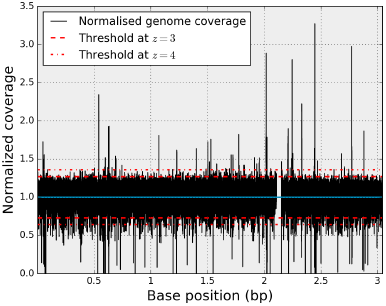
Normalized genome coverage 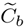 (bacteria test case). The outliers present in the original genome coverage *C_b_* (see Fig. 1) are still present as well as the deleted regions. The distribution is now centred around unity (blue line). Since the distribution is normalized, constant thresholds can be used (dashed lines). See section 3.2 for details.

### 3.2 Building a statistics

Since the reads are randomly generated (uniform distribution over the genome), when reads are mapped to the genome, the per-base coverage follows a Poisson distribution. It is discrete and has one parameter that corresponds to the sequencing depth (mean of the distribution). Yet, the Poisson distribution is often too narrow, as can be observed in the three test cases considered. This is due to biological over-dispersion. In order to account for over dispersion, the Poisson parameter can be distributed according to a second distribution. For instance when the Poisson parameter is distributed according to a Gamma distribution, we obtain a negative binomial, which has two shape parameters [Linder et al., 2013]. For large sequencing depth, we can approximate the negative binomial or Poisson distributions with a Gaussian distribution. We will use the mathematical notation 𝒩(*μ,σ^2^*) hereafter where *μ* is the average of the genome coverage (*δ* in an ideal case) and *σ* is its standard deviation.

Let us start with an ideal scenario where (i) there is no outliers, (ii) the running median window *W* is fixed and (iii) *δ*≫ 1. The latter means that *C_b_* distribution exhibits a Gaussian distribution ~ 𝒩(*μ,σ^2^*). Can we derive the distribution of the normalized genome coverage 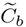 knowing that it is a ratio distribution? By definition, the numerator follows a𝒩(*μ,σ^2^*) distribution while the denominator’s distribution is the running median's distribution. The latter is generally not known, especially in the case of large *W*. Even if we knew the running median distribution, the ratio distribution is only known for two Gaussian distributions *X* and *Y* (Cauchy distribution) when (i) the two distributions are centred around zero, which is not the case, and (ii) when they are independent, which is also not the case. Further, the scenario we considered (no outliers, *W* fixed, *δ*≫ 1) is too restrictive since we are interested in identifying outliers and may encountered cases where *δ* is small (for which *C_b_* follows a negative binomial, not a Gaussian distribution).

Our first hypothesis is that 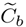 can be decomposed into a central distribution, 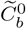, and a set of outliers,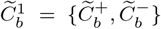where the central distribution is predominant: 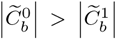, and where vertical bars indicate the cardinality of the sets.

Our second hypothesis is that the mixture model that represents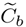 is a Gaussian mixture model of *k* = 2 models: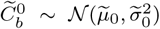 and 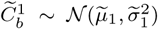. The Gaussianity hypothesis about the central distribution, 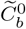 is valid as long as the raw sequencing depth is large (i.e., at least 10X). The Gaussianity of the outliers may be questioned, especially for the low-sampling case. However, in the context of a null hypothesis where the central distribution represents the background and the outliers the signal to detect, we can consider that the outliers population is a mix of samples and that we are in the limit of the central theorem. Similarly to the method deployed in [Linder et al., 2013] to identify a mixture model of negative binomials (on the raw genome coverage), we will use an Expectation Minimization (EM) [Dempster et al., 1977] method to estimate the parameters 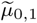 and 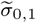 (on the normalised genome coverage).

The EM algorithm is an iterative method that alternates between two steps: (i) an Expectation step that creates a function for the expectation of the log-likelihood using the current estimate of the parameters, and (ii) a Minimization step that computes parameters maximizing the expected log-likelihood found in the first step. The likelihood function and the maximum likelihood estimate (MLE) can be derived analytically in the context of Gaussian distributions. Note that in addition to the means and standard deviations, the mixture parameters also need to be estimated. These are denoted 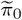 and 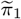. The EM algorithm is standard and can be found in various scientific libraries. Note, however, that the normalized genome coverage may contain zeros in the presence of deleted regions and the estimation of the mixture model should ignore them.

We have applied the EM algorithm on the normalized genome coverage vector on various real NGS data sets including the three test cases Fig. 4. The EM retrieves the parameters of the central distribution (in particular 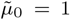) and the outliers. Note that the choice of the running median parameter,*W*, does not significantly affect the parameter estimation. In each case, the mean of the central distribution is very close to unity. The standard deviation varies significantly and is a function of the sequencing depth only (since the outliers are now incorporated in 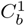. Finally, we can confirm that the proportion of outliers is small as compared to the central distributions by inspection of parameters *π*_0_ and 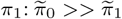.

**Fig. 4.**
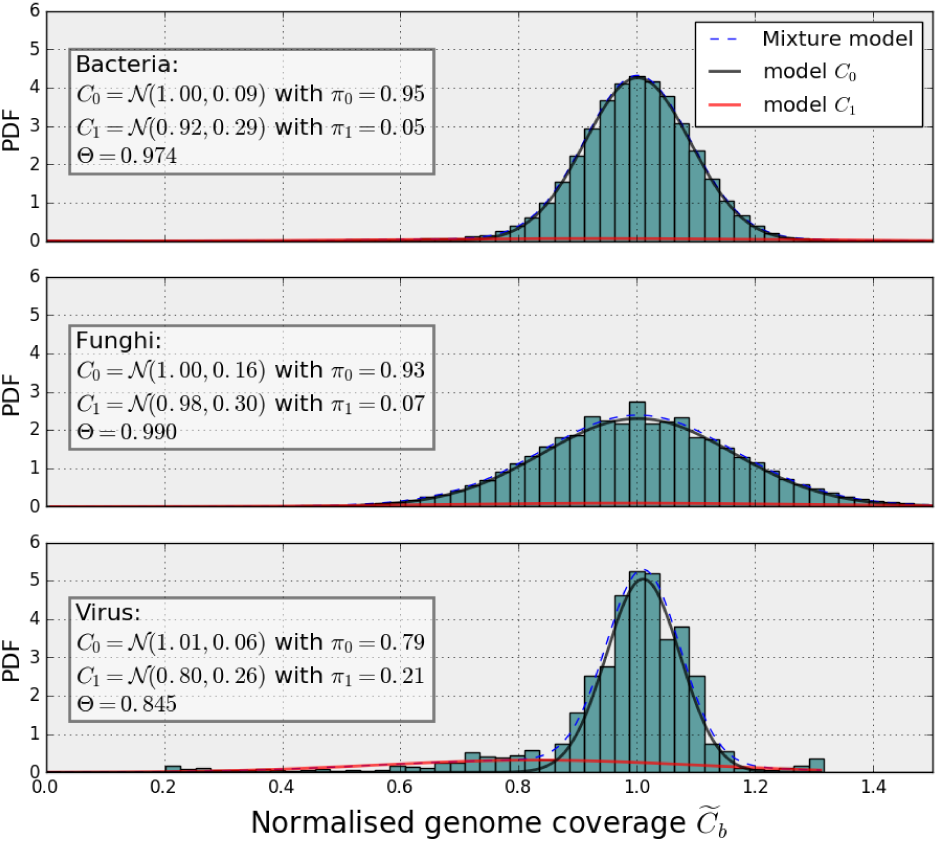
Probability density functions (PDFs) of the normalized genome coverage function concerning the three test cases. The distributions were fitted with a Gaussian mixture models with k = 2 models. The first model (black line) fits the central distribution’s PDF and the second model (red line close to y = 0) fits the outliers’ PDF. The dashed lines (close to the black lines) indicates the mixture distribution. In each panel, we report the parameters of the two Gaussian distributions, the proportions π_0_, π_1_ and the Θ parameter introduced in the text that gives the centralness of the data for each test cases.

Once we have identified the parameters of the central distribution 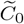, we can assign statistics for 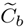 in terms of `z`-score:

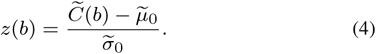

Since the `z`-score corresponds to a normal distribution, we can now set a threshold in terms of tolerance interval within which a specified proportion of the genome coverage falls. For instance, with a threshold of 3, we know from the normal distribution that 99.97% of the sample lies in the range −3 and +3. The exact mathematical value is given by the complementary error function, erfc(*x*), where 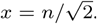 Note that for n = 3, 4 and 5, the tolerance interval is 99.73%, 99.993% and 99.999942%, respectively. Thus, for a genome of 1 Mbp, by pure chance we should obtain about 2700, 70 and 1 outlier(s), respectively.

If we now replace 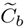 in Eq.4 using its expression from Eq. 3, we can express the original genome coverage as a function of the running median,the `z`-score and the parameters of the central distribution:

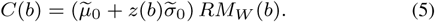

We can now set a fixed threshold *z(b)* = ±*n* in the normalized space. This is much easier to manipulate. Moreover, we can derive a variable threshold in the original space that is function of the genome position:

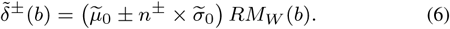

Examples of variable upper and lower threshold functions are shown in Fig. 1 and Fig.2 (red dashed lines). This manipulation results in a robust statistical estimate of the presence of outliers in the genome coverage. The `z`-score, computed earlier, provides a precise level of confidence.

Using the normalization presented above, we can define the **centralness** as one minus the proportion of outliers contained in the genome coverage:

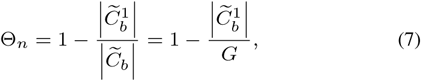

where *G* is the length of the genome, and vertical bars indicate the cardinality. This necessarily depends on how the threshold *n* is set in the normalized space. In the case of an ideal Gaussian distribution and *n* = 3, the centralness should equal the tolerance interval of a normal distribution 𝒩(0,1) that is the error function, erf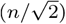. The centralness equals unity when there are no outliers i.e., *n* → ∞ to. Finally, note that the centralness is meaningless for values below 0.5 (meaning that the central distribution is not central!). As shown in Table.1, Θ_3_ equals 0.974, 0.99 and 0.86 in the three cases considered (bacteria, fungus, virus). So the proportion of outliers in the virus case is higher than in the two other test cases, which is not obvious at first glance given the very different lengths of the genome considered.

### 3.3 Genomic ROIs

Let us now consider the sub-set of outliers 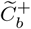. From the previous section, it is defined by positions that are above the fixed threshold *n*^+^ in the normalized space; it is a list of continuous or non-continuous positions; the list may be quite extensive for low threshold (e.g., for *n*^+^ = 2.5, the bacteria has 35 Kbp such positions). However, many positions belong the same event (*i.e.*, same cluster). Considering the short genome region in Fig. 5, which is made of 2000 base positions. It contains 5 different regions that cross the threshold *n*^+^. However, only one is well above. Ideally, the 5 events should be clustered together. To do so, we proceed with a double-threshold approach [Balasubramanian et al., 2005] where a second fixed threshold *m*^+^ is defined as *m*^+^ = *α*^+^ *n*^+^ where *α*^+^ ≤ 1 and usually set to 1/2.

**Fig. 5.**
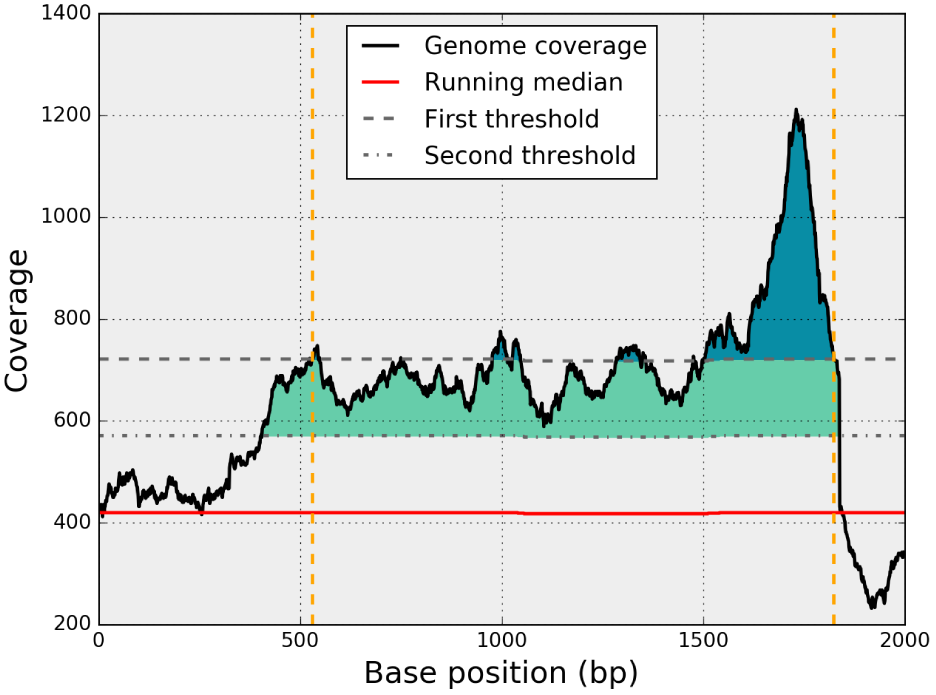
Example of a genomic region of interest (ROI) clustered using a double threshold method. The genome coverage (black line) and its running median (red) on a short genome location of 2kbp. The first threshold (top dashed gray line) alone identifies many short ROIs (dark blue areas). Using a second threshold (bottom dashed gray line), the short ROIs are clustered and identified as a single ROI (coloured areas). Yellow vertical lines indicates the beginning and end of the cluster.

In the normalized space, the double threshold method works as follows; We scan the entire genome coverage vector starting from the first position *b* = 0. As soon as a per-base coverage value crosses the threshold *m*^+^, a new cluster starts. We then accumulate following bases until the per-base coverage crosses *m+* again (going down). If the maximum of the cluster is above the first threshold, *n*^+^, then the cluster is classified as a region of interest. The process carries on until the end of the vector is reached. We repeat this classification for the lower case (with *m*^−^ = *α*^−^ *n*^−^). This method dramatically reduces the number of short ROIs. Finally, we can characterize each region with various metrics such as the length of the region, maximum coverage, mean coverage, mean and maximum `z`-scores.

## 4 Applications

Although the algorithm described is quite simple, each of the three steps need optimised implementation. We provide an implementation within the `Sequana` project [Desvillechabrol et al., 2016] as a standalone and as part of an original pipeline dedicated to variant calling. The standalone application is called `sequana_coverage` and is further described in the Appendix 6.2. Users can provide a BAM or a BED file [Quinlan, 2010] as input. Once the processing of the data is completed (running median, Gaussian mixture model, clustering of ROIs), a HTML report is created and users can browse the results. ROIs are provided as HTML tables that are downloadable as CSV files. If a reference is provided (it can be downloaded automatically via BioServices [Cokelaer etal., 2013] using `sequana_coverage ––download–reference <reference name>`), then the GC content of the reference versus the coverage along the reference is also calculated. With this material, users can use `sequana_coverage` in many different ways. Here is a non exhaustive list of applications.

**Table 1.**
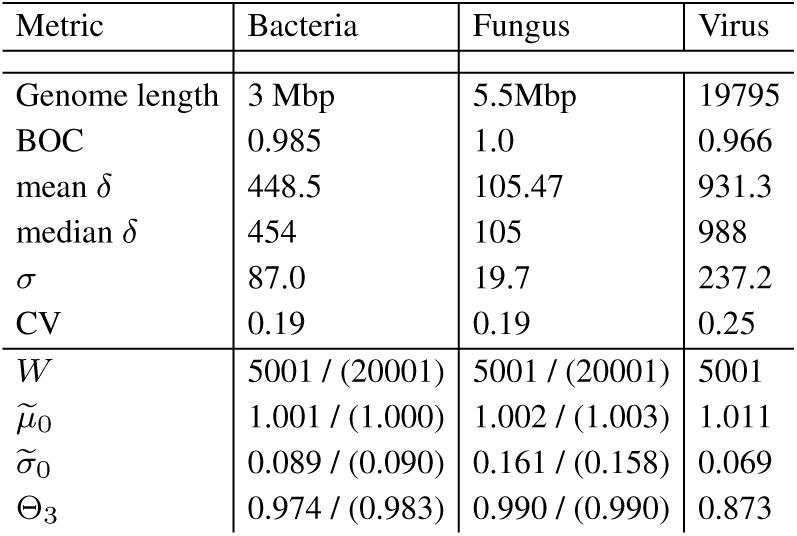
Metrics derived from the genome coverage of the three test cases considered (Bacteria, Fungus, Virus). The top part of the table contains metrics derived from the genome coverage only, while the bottom part contains metrics derived from the normalized genome coverage, 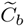. All metrics are defined in the text; BOC stands for breadth of coverage, *δ* for sequencing depth, CV for coefficient of variation. The standard deviation is denoted σ. In the bacteria and fungus cases, the running window *W* is set to 5 001 or 20 001 while for the virus we used 5 001 only. The parameters of the central distribution, 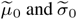 and the centralness, *Θ*_3_ are reported.

- Associate a statistic (z-score) on the genome coverage.
- Automatic detection of under or over covered ROIs. The CSV files provided may be used for further classification using machine learning tools. For instance the width of the ROIs and/or the maximum amplitude may be used as features.
- Effect of the GC content on the coverage using the reference. The GC content versus coverage plot is available in the report.
- Annotation of the ROIs if an annotated data file is provided (genbank). Again, it can be downloaded via the standalone application.
- Assess the quality of a *de-novo* remapping reads on contigs.
- Identify repeated regions. See hereafter for more details.

Regions of lower genome coverage are sometimes related to repeated content or unusual GC content [Dohm et al., 2008]. The identification of repeated regions is illustrated in Fig.6. We extracted specific regions where the filtered (mapping quality <30) and unfiltered coverage differ. Duplicated reads are removed in both conditions. The first difference occurs near a deleted region of 5kb around position 10,000. The second difference occurs around location 30,000. In the first case, we have deleted regions (the unfiltered coverage has zero coverage) compared to the reference. However, in the second case, the coverage of the unfiltered data seems normal with a value around 500. Only comparison with the filtered coverage allows us to identify that it is a repeated region. Moreover, using freebayes [Garrison, 2012] and the variant calling pipeline available (e.g., with the variant calling Sequana pipeline), no variants are found around this position. The report, and in particular the Javascript plots, allows us to identify all such repeated regions. In the case of the bacteria used in this example, looking at the 3Mbp genome, about 70 repeated regions with at least 50 bases and an average difference in coverage of 50X can be identified.

**Fig. 6.**
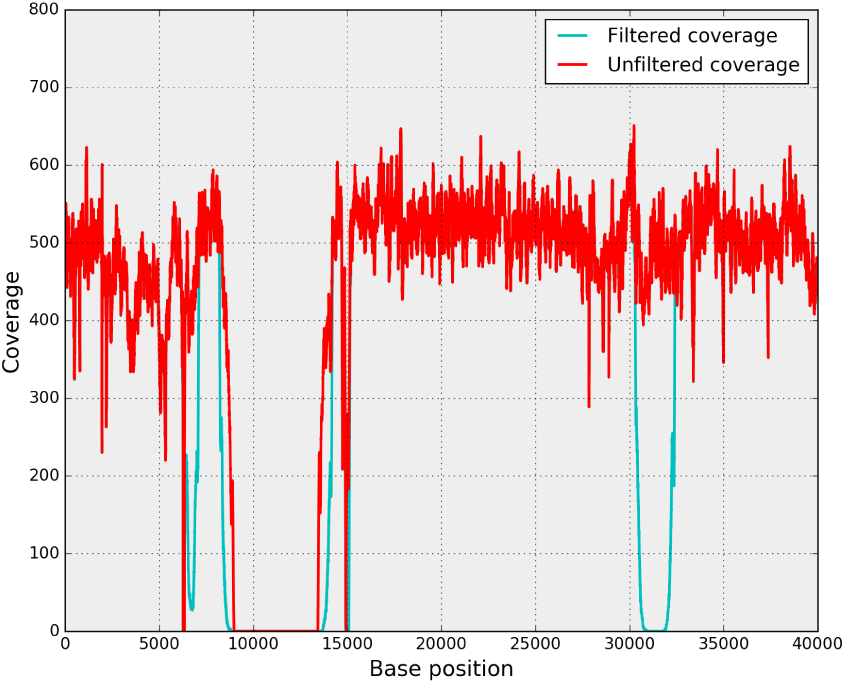
Identification of repeated regions by comparing the genome coverage before and after filtering of low quality mapped reads. In the filtered coverage series (blue), the mapped reads with low quality were removed. In particular the duplicated reads are removed. This allows us to identify repeated regions easily.

## 5 Conclusion

The genome coverage along a reference contains valuable information and deserves to be part of an NGS toolkit (e.g., quality control of an alignment before a variant detection pipeline). Yet, it is too often summarised by its sequencing depth even though the raw data usually contains a wide spectrum of features such as deleted regions, low frequency trends, nonhomogeneous central distribution, repeated regions, …

The method presented in this paper provides a robust statistical framework to detect under and over-covered genomic regions that can be further characterized with basic statistics (length, mean coverage, maximum `z`-score, …). Although robust, the method remains simple and can be summarized in three main steps: (1) detrending of genome coverage series using a running median (ii) parameter estimation of the central distribution of the normalized genome coverage series using an EM approach (for a Gaussian mixture model), (iii) clustering and characterization of the outliers as genomic regions of interest (ROI).

We underlined the value of the running median algorithm as compared to a moving average while emphasizing the practical impact of the running median algorithm complexity. Indeed, an efficient implementation is of paramount importance in the context of NGS analysis. In addition, circular series and multi-chromosome organisms should be handled. We wrap the algorithm within the standalone application `sequana_coverage`, which also provides HTML reports with a summary of the genomic regions of interest. The HTML reports provide visual introspection of the genome coverage, list of geomic ROIs and statistics such as the centralness, a metric that encompasses the preponderance of the central distribution with respect to the outliers.

Although we presented test cases with relatively large sequencing depth (100X to a thousand), it is based on a robust statistics and practical cases down to 30X were studied with success. We believe that the algorithm can be used to sequencing depth as low as 10X. Below 10X, a Gaussian distribution hypothesises not valid and the `z`-score values are less precise. A natural extension to this work is to consider low sequencing depths below 10X

With additional features such as the ability to annotate the ROIs with genbank files and the identification of repeated regions, we believe that the standalone application `sequana_coverage` will help researchers in deciphering the information contained in the genome coverage.

## Acknowledgement

We are grateful to Nicolas Escriou (Institut Pasteur) for providing the FastQ and reference of the Virus test case. We are also grateful to Benoit Arcangioli (Institut Pasteur) and Serge Gangloff (Institut Pasteur) for providing the FastQ files and reference of the *S. Pombe* test case.

## 6 Appendix

### 6.1 Running median implementation

The mean is a measure of the central tendency of a population. It is not a robust estimator in the presence of large extraneous outliers in the population. In such a situation, it is preferable to consider a truncated mean or a *median* estimator. The median is the middle point of a sample set in which half the numbers are above the median and half are below. More formally, let us consider a sample *s[i]*, *i* = 1,…, *n* and *S[i]* the sequence obtained by sorting *s[i]* in ascending order (ordering of equal elements is not important here). Then, the median is defined as

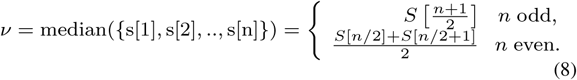

Let us now consider a series *X*(*k*) where *k* = 1,…,*N*. Then, the *running median* of X(k) is defined as the sequence *v*(*k*) = median({X(k),X(k + 1),…,X(k + W)}), *k* = *W*/2,…,*N* – *W*/2 where *W* is a window size defined by the user and the application. The first *W*/2 and last *W*/2 values are undefined so we should have *W ≪ N*.

Since we perform a sorting of an array of *W* elements at *N* positions, the complexity of the running median is *N* times the complexity of the sorting algorithm. If *W* and *N* are small (e.g., removal of narrow lines in power spectral density in addition to the overall smoothing of time or frequency series [Balasubramanian et al., 2005]), a naive quick-sort algorithm (𝒪(*W^2^*) in the worst case scenario) may be used. However, better algorithms do exist and can be decreased to 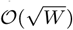 in the worst case as implemented in [Mohanty, 2002]. Yet, in NGS applications, *N* could easily reach several millions and *W* may need to be set to large values up to 50,000 (e.g., to identify long deleted regions).

Instead of computing the median at each position, *k*, a more efficient solution consists in re-using the sorted block at *k −* 1, and to maintain the block sorted as new elements are added. Indeed, one only needs to *insert* the next sample into the sorted block and *delete* the earliest sample from the sorted block. A standard Python module named *bisect* provides an efficient insertion in sorted data (keeping the data sorted). The complexity of this sorting algorithm is 𝒪(log *W*).

So far, we have neglected the cost of the insertion and deletion steps, which is not negligible. For instance, in Python language, one of the most common data structure is the *list*. It is a dynamically-sized array (i.e., insertion and deletion of an item from the beginning or middle of the list requires to move most of the list in memory) and the look-up, insertion and deletion have a 𝒪(*n*) complexity. So the running median is actually dominated by the slow 𝒪(*n*) insertion and deletion steps. A better data structure is available thanks to the *blist* package; it is based on a so-called B-tree, which is a self-balancing tree data structure that keeps data sorted. The blist allows searches, sequential access, insertions, and deletions in 𝒪(log *n*) (see https://pypi.python.org/pypi/blist/ for details).

Based on materials from http://code.activestate.com/recipes/576930/, we have implemented a running median function in Python within **Sequana** [Desvillechabrol et al., 2016]. A simplified version of which is reproduced here below. It is fully described and documented in the official on line documentation at http://sequana.readthedocs.org

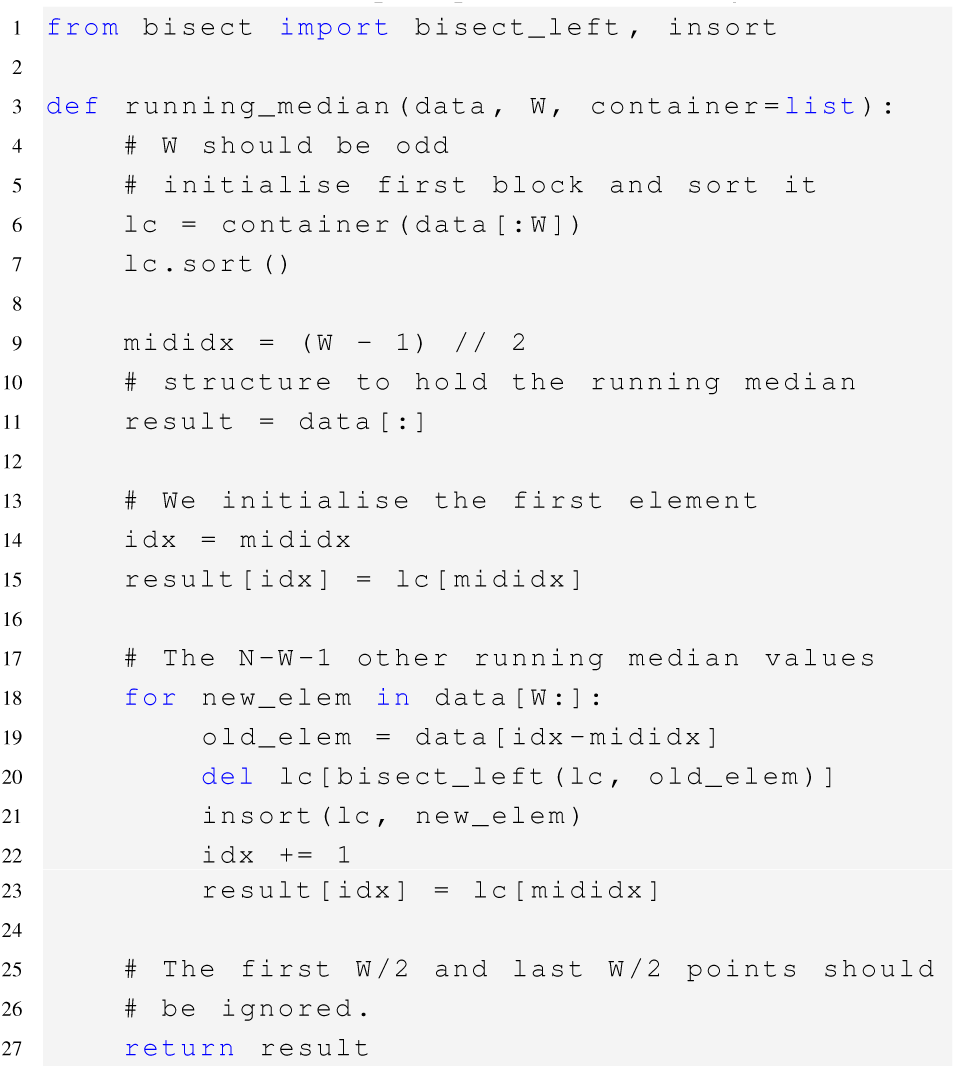

In Fig.7, we compare the performance of two variants of our implementation and the running median implementation available within `SciPy` [Jones et al., 2001]. For *W* > 20, 000 up to 200,000, our implementation is 2–3 order of magnitude faster than the SciPy version. We should emphasize the fact that the `SciPy` function has additional features since it is available for **N**-dimensional data sets whereas we restrict ourselves to 1-D data sets. In `Sequana`, the two variants only differ in the data structure being used to hold the data (list versus `blist`). The Fig. 7 shows the difference between the list and `blist` data structures that is marginal for low *W* values while for large values asymptotic behaviours are reached showing the interest of the blist over the list choice.

**Fig. 7.**
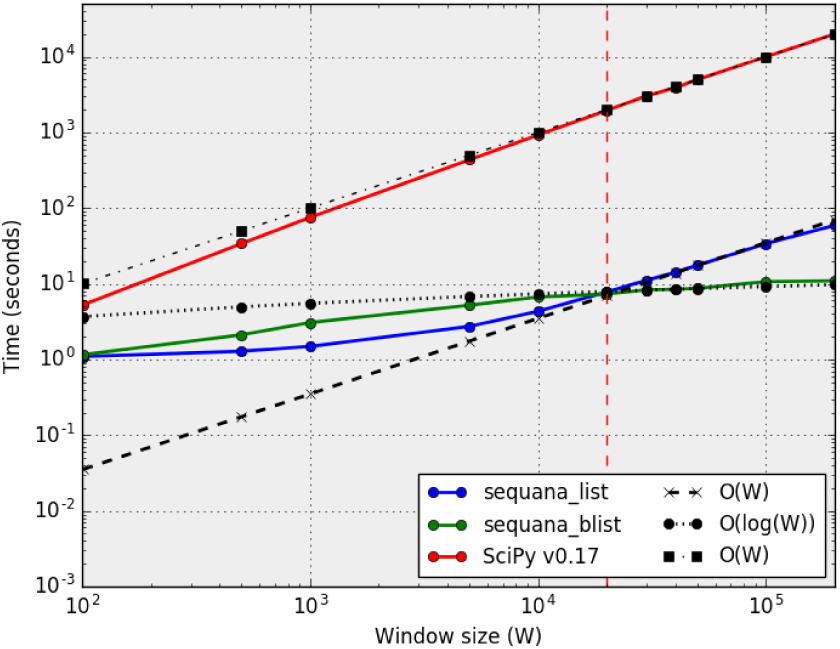
Computational cost of running median algorithms as a function of the window size parameter W (for *N* = 1*e*6). Three variants are compared: `SciPy` [Jones et al.,2001] implementation (function medfilt v0.17) and the `Sequana` implementations using the `list` and `blist` data containers (see text for details). The `SciPy` variant has a𝒪(*W*) complexity irrespective of the *W* value. For low *W* values (*W* > 20; 000), the two Sequana variants have 𝒪(*log*(*W*)) complexity. For larger *W* values, the blist keeps its 𝒪(*log*(*W*)) complexity while the list container follows a 𝒪(*W*) complexity. The computational time of the Sequana implementations are about 2 orders of magnitude below the `SciPy` one.

### 6.2 Standalone application

`Sequana` is a Python library that provides NGS pipelines in the form of *snakefiles* based on the workflow management system called Snakemake [Köster and Rahman, 2012] (Makefile-like with a Python syntax). `Sequana` also provides a Python library with re-usable blocks. Moreover, we provide independent standalone applications. One of them is called `sequana_coverage`; it includes the different features related to genome coverage exposed in this paper. Although the standalone `sequana_coverage` has a self-explanatory help, we give here below an example that shows how to generate an HTML report from a BED file. The BED being a data structure that stores the genome coverage information [Quinlan, 2010] (3-columns tab-delimited file). A basic usage is as follows:

`sequana_coverage ––input virus.bed −w 4001 −o`

Several chromosomes may be present (e.g., fungus case). By default, the first chromosome is used but one can provide the chromosome number using the *−c* option. The *−o* option indicates that the input is made of a circular DNA. The running median window can be tuned using *-w* option. Full details are available using *−−help*. An HTML report is created by default in the .`/report` directory. In the case of long genome (larger than 0.5 Mbp), independent `JavaScript` pages are created to focus on 0.5 Mbp-long regions to speed up browsing and introspection. A list of genomic regions are available as HTML tables but also as downloadable CSV files. An additional feature is the ability to download a reference genome (given its ENA or NCBI accession number). This is achieved internally using BioServices [Cokelaer et al., 2013] that can switch between the ENA or NCBI web services to download the data programmatically. This is particularly useful to further compare the genome coverage with e.g., the GC content of the reference. Finally, let us note that the standalone application is scalable: the virus case takes a few seconds while the 5Mbp bacteria case takes about one minute on a standard computer including analysis and HTML reports (Python implementation). An HTML report example is available on http://sequana.readthedocs.org. A docker file available via the documentation should help users experimenting with their own data set.

### 6.3 Terminology

The terms *depth* and *coverage* have been used interchangeably in many papers. This is especially the case for the *coverage* term that is a short form for different notions such as breadth of coverage, depth of coverage, genome coverage and so on. In the seminal work from [Lander and Waterman, 1988], the authors introduced the theoretical notion of **redundancy of coverage**. This is well defined as *N_r_ L_r_* /*G*: the number of experimental reads, *N_r_*, times the fixed length of the reads, *L_r_*, divided by the genome length, *G*. Note that it is a metric derived from the raw data, not the mapped reads. Following this work, many synonyms have been introduced: **sequencing depth** [Sims et al., 2014], **Depth of coverage** (DOC), **fold-coverage**, or simply, *depth*, or *coverage*. The two later terms being ambiguous short forms for depth of coverage. Since, *coverage* may be used as a short form for depth of *coverage* but also genome coverage, which are two different notions, we do not recommend the use of short forms.

Instead, we define the **genome coverage** as the number of reads mapped to a specific position within a reference genome. It is a function of the position on the genome of reference. From the genome coverage we derive the **sequencing depth**, which is the average of the genome coverage.

For completeness, note also the term **breadth of coverage** (BOC): the assembly size divided by the target size. It can also be the proportion of total intended genome representation in the data set with at least some sequencing depth.

